# Single-molecule kinetics of pore assembly by the membrane attack complex

**DOI:** 10.1101/472274

**Authors:** Edward S. Parsons, George J. Stanley, Alice L. B. Pyne, Adrian W. Hodel, Adrian P. Nievergelt, Anaïs Menny, Alexander R. Yon, Ashlea Rowley, Ralf P. Richter, Georg E. Fantner, Doryen Bubeck, Bart W. Hoogenboom

**Affiliations:** London Centre for Nanotechnology, University College London, London WC1H 0AH, United Kingdom; Institute of Structural and Molecular Biology, University College London, London WC1E 6BT, United Kingdom; Laboratory for Bio- and Nano-Instrumentation, Swiss Federal Institute of Technology Lausanne (EPFL), 1015 Lausanne, Switzerland; Department of Life Sciences, Imperial College London, South Kensington Campus, London SW7 2AZ, United Kingdom; School of Biomedical Sciences, Faculty of Biological Sciences, University of Leeds, Leeds LS2 9JT, United Kingdom; School of Physics and Astronomy, Faculty of Mathematics and Physical Sciences, University of Leeds, Leeds LS2 9JT, United Kingdom; Astbury Centre for Structural Molecular Biology, University of Leeds, Leeds LS2 9JT, United Kingdom; Department of Physics and Astronomy, University College London, London WC1E 6BT, United Kingdom

## Abstract

The membrane attack complex (MAC) is a hetero-oligomeric protein assembly that kills pathogens by perforating their cell envelopes. The MAC is formed by sequential assembly of soluble complement proteins C5b, C6, C7, C8 and C9, but little is known about the rate-limiting steps in this process. Here, we use rapid atomic force microscopy (AFM) imaging to show that MAC proteins oligomerize within the membrane, unlike structurally homologous bacterial pore-forming toxins. C5b6 interacts with the lipid bilayer prior to recruiting C7 and C8. We discover that incorporation of the first C9 is the kinetic bottleneck of MAC formation, after which rapid C9 oligomerization completes the pore. This defines the kinetic basis for MAC assembly and provides insight into how human cells are protected from bystander damage by the cell surface receptor CD59, which is offered a maximum temporal window to halt the assembly at the point of C9 insertion.

## Introduction

The formation of lethal membrane pores is a ubiquitous event in defence and attack between pathogens and their hosts^1-3^, It plays a critical role in the lytic, antimicrobial activity of human serum, the discovery of which was an early milestone in immunology^4^, and decades of study have unravelled the interplay between proteins that lead to the formation of immune pores in microbial membranes. The formation of these membrane attack complex (MAC) pores represents the final step in the activation of the complement system, an integral component of innate immunity, which surveys our body for pathogenic bacteria and which ‘complements’ the ability of leukocytes to kill pathogens. Dysregulation of MAC formation has been implicated in human disease^5,6^, and therapeutics that control complement are being harnessed for cancer immunotherapy^7,8^. Understanding how complement proteins assemble from innocuous soluble monomers into killer transmembrane pores can therefore contribute to developing strategies for treating human disease where the MAC is implicated^5^, and for repurposing the complement system as a potent immunotherapeutic^9^.

Assembly of the MAC is the end product of a complex series of biochemical interactions in which initially soluble complement proteins bind and undergo dramatic structural rearrangements to form a transmembrane pore. The resulting MAC pore is a hetero-oligomer formed from the irreversible, stepwise assembly of 7 different polypeptide chains: C5b, C6, C7, C8 (a hetero-trimer comprised of C8α, C8β and C8γ) and C9, where 18 copies of C9 are required to complete the pore (***Fig. 1a***, inset). Triggered upon detection of a pathogen, activation of complement leads to the generation of C5b via the cleavage of C5 by membrane-bound C5-convertase enzymes^10^. C5b is a metastable intermediate that rapidly sequesters C6^11^. Recruitment of C7 unfurls a lipophilic domain upon binding, while integration of C8 into the assembly is accompanied by an initial insertion into the membrane. The C5b-8 initiator complex then binds C9 and undergoes unidirectional oligomerization (with 18 copies of C9) to complete an 11 nm wide transmembrane pore, as characterized in increasing structural detail by cryo-electron microscopy (cryoEM)^12-16^. Together with crystallographic structures of component proteins, high-resolution cryoEM analyses of the full pore have identified regulatory roles for auxiliary domains^15,17,18^ that control the transition from stable proteins in our blood to lethal transmembrane pores. However, it remains unclear which are the rate-limiting steps in the assembly pathway of these complement proteins.

**Figure 1.**
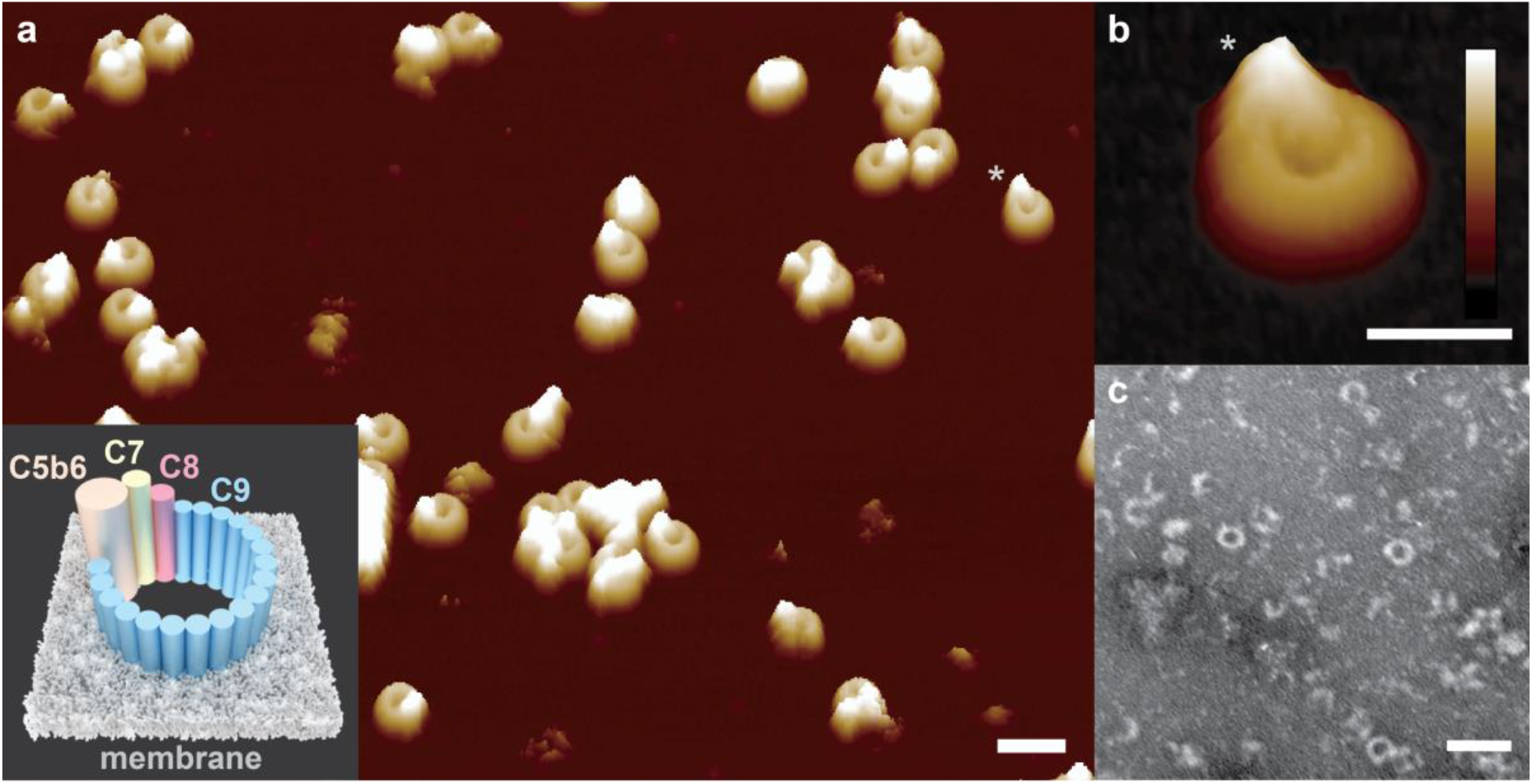
Formation of MAC on bacterial model membranes. **a,** 3D AFM representation of the endpoint MAC pore structure on supported bilayers composed of an *E. coli* lipid extract. Inset: Schematic of the MAC, self-assembled from complement proteins C5b6, C7, C8 and C9, embedded within a lipid membrane. **b,** Zoom-in of a single MAC pore (marked with an asterisk in a). **c,** Negative-stain EM of *E. coli* lipid bilayers deposited on silicon dioxide grids, sequentially incubated with complement proteins C5b6, C7, C8 and C9, resulting in characteristic MAC rings observed in the bilayer membrane. Scale bars: a,c: 50 nm, b: 25 nm. Height scale (scale inset in b), a,b: 20 nm.

As there is no known lipid or receptor specificity for MAC membrane insertion, the kinetics of pore assembly drives both the rapid innate immune response to pathogens and dictates how the MAC can be most effectively inhibited on membranes of self-cells. For human cells, the only known membrane-associated inhibitor of MAC assembly is CD59, a glycosylphosphatidylinositol (GPI) anchored cell surface receptor. CD59 binds the transmembrane residues of C8 and C9^19^, preventing pore formation and further oligomerization of C9^20^. The kinetics of MAC formation must allow a temporal window such that inhibitory factors can interfere at appropriate stages in the assembly pathway. Therefore, a kinetic analysis of MAC assembly will provide a much-needed framework to understand how CD59 inhibits lysis.

To understand the molecular mechanism and kinetics underpinning how and when the MAC assembly becomes cytolytic, we sought to track the progression of the complement terminal pathway at the level of single pores. Using rapid atomic force microscopy (AFM) imaging on supported model membranes, we visualize the initial interactions of complement proteins with the membrane, and resolve the kinetics of MAC pore formation. Together these data reveal the overall rate of the assembly process and identify which steps in the pathway are rate-limiting.

## Results

### MAC forms pores in bacterial model membranes

To enable AFM tracking of MAC self-assembly at single-molecule resolution, we developed a model membrane system that supported the formation of transmembrane pores. We sequentially incubated complement proteins C5b6, C7, C8 and C9 at physiological concentrations^10,21^ on supported bilayers formed from *E. coli* lipid extract^22^ and from *pseudo E. coli* lipid mixtures comprised of 1,2-dioleoyl-sn-glycero-3-phosphocholine (DOPC), 1,2-dioleoyl-sn-glycero-3-phospho-(1’-rac-glycerol) (DOPG) and 1,2-dioleoyl-sn-glycero-3-phosphoethanolamine (DOPE). In our model comprised of synthetic lipids, we sought to harness the physico-chemical properties of the bacterial membrane: PG lipids harbour negative charge in their phosphoglycerol headgroup, whilst PE introduces a degree of stored curvature elastic stress into the plane of the membrane^23^. Inspection by AFM and fluorescence recovery after photobleaching (FRAP) reveals a single continuous supported bilayer with rapid in-plane diffusion (***SI Fig.*** 1).

High-resolution AFM images of the resulting end-point MAC pores are consistent with cryo-EM reconstructions *(**SI Fig. 2a-d**)*. We clearly resolve the lumen of the β-barrel pore in addition to the protruding C5b stalk that hallmarks the MAC *(**Fig. 1**)* ^11,13^. Vesicle lysis assays corroborate the functional requirement of both a C5b-8 ‘initiator’ and C9 ‘propagators’ for MAC to form lytic pores (***SI Fig. 2e***); and negative-stain EM on lipid bilayers shows that C5b-8 is required for the oligomerization and insertion of C9 within the membrane (***SI Fig. 3***). Together, these data confirm that the C5b-8 initiator complex, comprised of C5b6 in complex with C7 and C8, is essential for the formation of a functional MAC within the bacterial model membrane.

### C5b6 binds to bacterial target membranes to initiate MAC assembly

Previous biochemical studies of MAC formation were performed on an ensemble of erythrocyte and liposome membranes, preventing analysis of individual pores^24-26^. By contrast, our experimental system facilitates a stepwise study of MAC assembly at the single-molecule level. Upon addition of C5b6 to a bacterial model membrane, we observe features of few-nanometre dimensions *(**Fig. 2a**)* that appear at different locations in subsequent images. Their mobile nature on the membrane and protruding structure pose a challenge for AFM imaging^27,28^, in practice putting them at the detection limit of AFM. Further *in situ* incubation with C7 yields an increase in the number of such features, which appear at enhanced clarity and robustness after the subsequent addition of C8 *(**Fig. 2a, SI Fig. 4***). These data indicate that C5b6 binds to the bacterial model membrane and becomes more static and/or more robust against the movement of the scanning AFM tip following assembly with C7 and C8.

**Figure 2.**
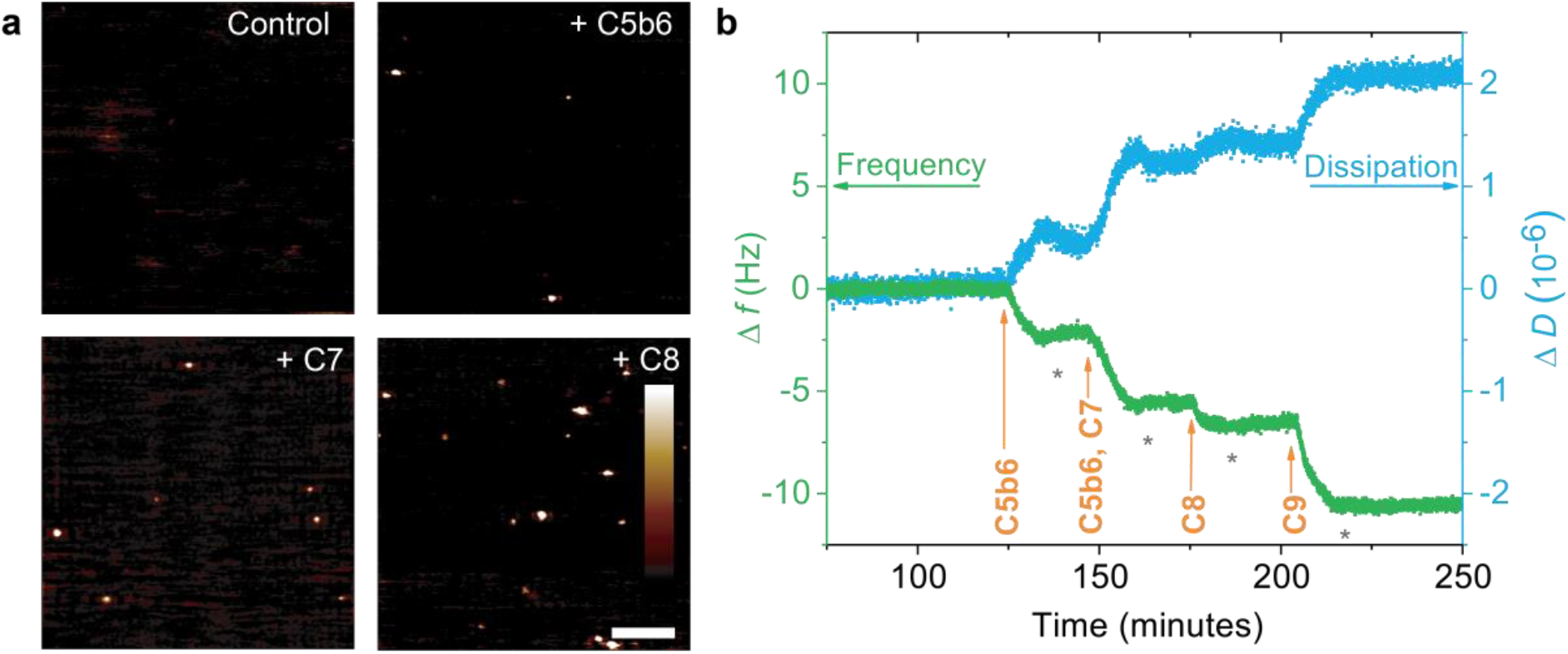
C5b6 initiates MAC formation by binding to the bacterial membrane. Binding of complement components C5b6, C7 and C8 to supported bacterial model membranes containing PG lipids, as observed by AFM and QCM-D. **a,** AFM images increasingly show protruding features upon addition of C5b6, C7 and C8 to a bilayer formed of DOPE:DOPG (50:50 mol %). Scale bar: 200 nm, height scale: 1 nm. **b,** QCM-D shows binding as distinct steps in both frequency (Δ*f*) and dissipation (Δ*D*) upon sequential addition of C5b6, C7, C8 and C9 to a lipid bilayer formed of DOPC:DOPE:DOPG (47.5:47.5:5 mol %) on a silicon dioxide QCM-D sensor; wash steps (*) between additions do not lead to a reduction in the signal size, revealing that the protein binding to the membrane is stable. The dissipation shift increases as protein addition augments the softness of the lipid bilayer film.

To rule out artefacts due to the inherent invasiveness of the AFM measurement, we also performed binding assays by quartz crystal microbalance with dissipation monitoring (QCM-D), which correlate with our AFM data. Briefly, we prepared supported lipid bilayers on a silicon dioxide QCM-D sensor, with which we could detect membrane binding of complement proteins^29,30^ (***Fig. 2b, SI Fig 5***). Upon addition of C5b6, we observe shifts in frequency (Δ*f*) and dissipation (Δ*D*) that persist upon washing with buffer. These data confirm C5b6 binding to the membrane and demonstrate the stability of this membrane-bound state. Subsequent incubations with C7 and C8 show further stable complexes that again could not be removed by washing with excess buffer. Upon addition of C9, we observe a final shift in frequency, corresponding to the inserted MAC pore. Taken together, these results demonstrate that the MAC is membrane-bound throughout its entire assembly pathway, beginning with C5b6.

### C9 monomers are recruited directly from solution

MAC proteins are structurally homologous to the immune protein perforin and to bacterial cholesterol dependent cytolysins, both of which form pores through the homo-oligomerization of membrane-associated monomers.^3,27^ We next explored whether the hetero-oligomeric MAC pore could assemble via analogous membrane-bound C9 oligomers, prior to its association with C5b-8. However, from the absence of C9 upon *E. coli* lipid bilayers in the electron microscopy and AFM images in ***SI Fig. 3***, we conclude that such intermediates, if existing at all, are only transiently bound to the membrane. Upon addition of C9, AFM imaging (***SI Video 1***) does not show the characteristic pre-pore carpet that was detected by AFM for perforin^27^ and the cholesterol dependent cytolysin suilysin^28^. Instead, we observe a planar membrane background that remains featureless until MAC pores emerge, implying the absence of any pre-pore oligomers until the pore is assembled within the membrane. The static nature of these MACs confirms their membrane-inserted, pore character^27^. Taken together, these data indictate that C9 monomers are recruited directly from solution to the nascent pore and not via a membrane-bound C9 intermediate.

### Initial insertion of C9 is a kinetic bottleneck in MAC assembly

By tracking the appearance and evolution of individual pores, we next determined the reaction kinetics that govern MAC assembly. Upon association with C5b-8, C9 oligomerizes to complete a transmembrane pore. With a frame rate of 6.5 sec per frame, we used AFM to visualize C9 oligomerization in real time at 30 °C (***Fig. 3a, SI Video 1***, in which *t* = 0 is defined as the time of C9 addition); data recorded at the physiological 37 °C showed similar kinetics (***SI Fig. 6, SI Video 2***). A single MAC pore is formed within the first few frames immediately after addition of C9. The pore persists in isolation for 30 seconds, after which multiple (3-5) pores appear simultaneously, and the number of pores augments at a gradually decreasing rate, up to approximately 50 MAC pores in the field of view at the end of the recording (***SI Video 1*** and ***Fig. 3a***).

**Fig. 3:**
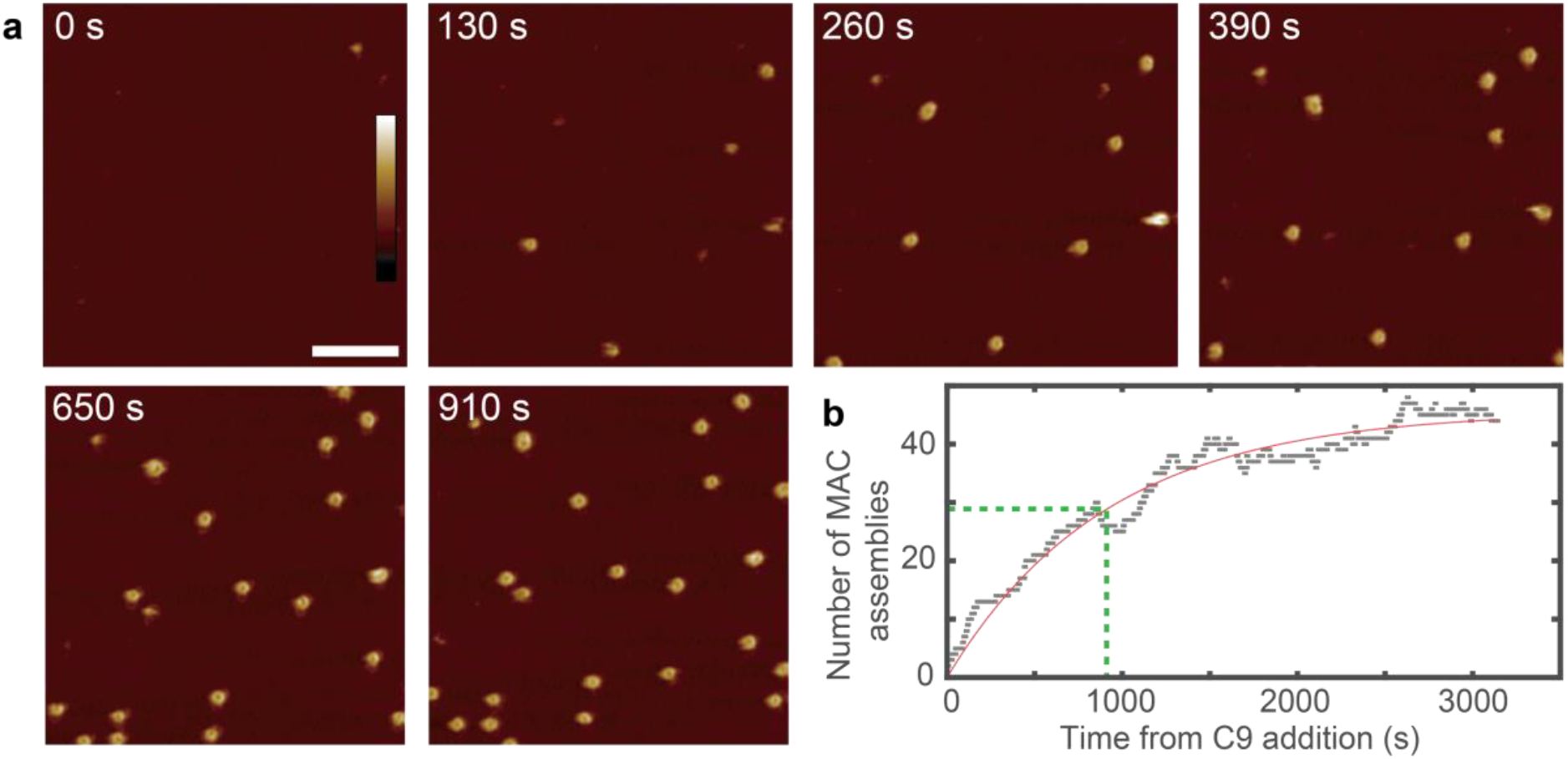
AFM shows that initial insertion of C9 is rate limiting in MAC assembly. **a,** Time-lapse AFM imaging of MAC assembly on supported bilayers formed of *E. coli* lipid extract (see ***SI Video 1*** for full data set). MAC assembly events initiate at distinct time points, with each MAC pore rapidly completing within a few frames; this implies a rate-limiting-step corresponding to initial insertion of C9. Scale bar: 200 nm; height scale (colour scale bar inset in image for *t* = 0 min): 40 nm. **b,** Number of MAC assemblies detected (see Methods) with time, indicated as grey squares; the solid red line represents fitting with the function *A* (1 – exp(–*t*/*τ*_init_)), where *τ*_init_ = 912 ± 32 s represents the characteristic time of MAC pore appearance, highlighted by the dashed green lines. The completion time of each individual MAC is much shorter than *τ*_init_.

Remarkably, complete MAC pores continue to appear over an hour; yet the completion of each individual MAC occurs at the timescale of a few seconds to minute (***Fig. 3a, SI Video 1***). Hence, two distinct kinetic steps are observed upon C9 addition to the nascent MAC. Firstly, we observe slow C9 binding to C5b-8, which is taken as our initial detection of a pore-forming event after the addition of C9. We define kinetics of this event by a characteristic initiation time τ_init_. Secondly, our data show rapid C9 oligomerization to a C5b-8C9n MAC pore. We define the time for each growing pore to fully assemble as τ_olig_. Consistent with the description of these two distinct kinetic steps, the vast majority of end-point MACs in our data are complete, ring-shaped pores. If the rate of initial C9 insertion were faster than the subsequent oligomerization reaction, kinetically-trapped, arc-shaped assemblies would occur due to monomer depletion, as has been observed for other pore-forming proteins^27,28,31^. To determine whether the kinetic bottleneck is attributed to C8 incorporation or insertion of the first C9, we allowed an extended temporal window after the addition of C8 and prior to adding C9 (***SI Fig. 7, SI Video 3***): we confirm by time-lapse AFM imaging that this does not lead to more efficient/faster MAC formation (***SI Fig. 7, SI Video 3***).

### Rapid AFM imaging allows quantification of reaction times

To quantify the reaction time of initiation and oligomerization, we analysed rapid AFM imaging data documenting pore assembly. Considering the nearly 100-fold difference between the timescales of MAC appearance (initiation) and completion (C9 oligomerisation), the timescale of completion can be assumed negligible relative to that of initiation. We detected pores by cross-correlation during particle tracking (see methods), and take pore appearance as a proxy that reports on the time of initiation of MAC assembly after addition of C9 to the reservoir. In doing so, we determine a characteristic initiation time τ_init_ = 912 ± 32 s (***Fig. 3b***).

The kinetics of C9 oligomerization were determined by tracking areas with individual pores from just before the point of initial detection to after completion (***Fig. 4, SI Video 4***). Image sequences of these tracks show distinct intermediates of a growing pore as the MAC completes (***Fig. 4a, SI Fig. 8***). To quantify the timescale of the transition we use the average frame height (defined by the average pixel intensity) as a proxy to report on the completeness of the pore. As the pore evolves we observe a gradual increase in the average height in the frame, which plateaus as the MAC reaches a final state (***Fig. 4b, SI Fig. 9***). Previous structural studies defined the stoichiometry of the complete MAC as having 18 copies of C9^12^. To measure the time required to add 17 copies of C9 after the initiation event, *τ*_olig_, we record the width of the transition between detection of the initial event and appearance of a complete MAC pore, *τ*_olig_ = 112 ± 17 s (***Fig. 4c***). This is an order of magnitude shorter than *τ*_init_, and implies that the average time per addition of each of the remaining 17 C9 subunits (*τ*_+_ = *τ*_olig_/17 = 6.6 ± 1.0 s) is more than two orders of magnitude shorter than *τ*_init_.

**Fig. 4:**
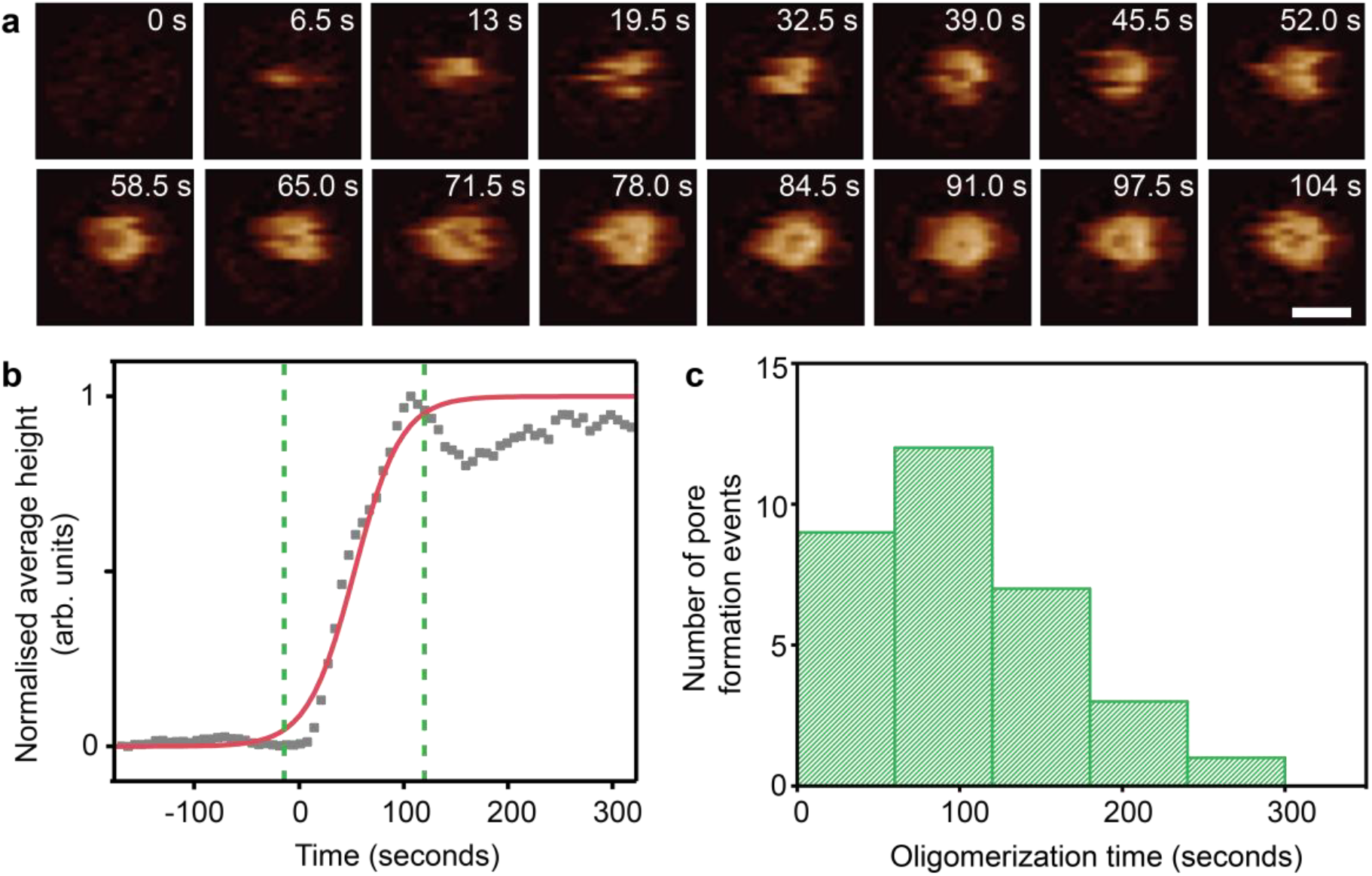
Real-time imaging of C9 oligomerization. **a,** AFM image sequence of MAC assembly, cropped from data shown in ***Fig. 3*** and ***SI Video 1***. C9 oligomerization completes within frames shown (104 s, 6.5 s/frame; 0 s is here approximately defined by the frame preceding detection of the growing MAC). Scale bar: 30 nm, height scale (see colour scale bar in ***Fig. 1***): 16 nm. **b,** The normalized average frame height versus time for a single pore forming event (corresponding to event shown in a), plotted here as a measure for completion of MAC assembly. The red line represents a sigmoidal fit to the data, as a generic and mathematically convenient description of a smooth transition between pore absence and pore completion. The C9 oligomerization time is determined from the width of the transition, highlighted by green dashed lines (see ***SI Fig. 9*** for details). **c,** Distribution of oligomerization times, extracted from *n* = 33 isolated pore forming events in 6 independent experiments.

### A kinetic model for C9 assembly in the MAC

Given that *τ*_init_ ≫ *τ*_+_, we can interpret our data in terms of the separate reactions C5b-8 + C9 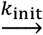 C5b-8C9 and 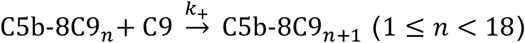, with respective rate constants *k*_init_ and *k*_+_. We observe that the C9 oligomerization times are independent of the time at which the reaction is initiated (***SI Fig. 10***), consistent with C9 being in excess. As such, the interpretation of this reaction scheme can be further simplified by the assumption of an excess and thus approximately constant C9 concentration in solution ([C9] ≈ 1.4 μM) over the duration of our experiments. Consequently, under our experimental conditions the final number of MAC pores is set by the number of C5b-8 complexes on the membrane. These assumptions lead to the exponential time dependence of MAC appearance as observed in ***Fig. 3b***, with *k*_init_ ≈ *τ*_init_^-1^[C9]^-1^ = 0.78 s^-1^mM^-1^ (see Methods). In addition, assuming a constant rate of C9 addition after the insertion of the first C9, we find *k*_+_ ≈ *τ*_+_^-1^[C9]^-1^ = 108 s^-1^mM^-1^ (see Methods). This more quantitative analysis confirms that the initial insertion of C9, together with its binding to C5b-8, is the major rate-limiting step in MAC assembly. Interestingly, this rate-limiting step coincides with the stage where MAC pore formation is inhibited by CD59, which is present on the surface of human cells to prevent them from being lysed by complement^20,32^.

## Discussion

The MAC represents a biomedically important system in which to probe how unique individual proteins self-assemble into a macromolecular functional unit. Once initiated, five soluble complement proteins sequentially and irreversibly self-assemble into a hetero-oligomeric pore that opens up an 11 nm hole in a fluid lipid bilayer. While there is an extensive (and still expanding) body of structural and functional information documenting the MAC and its constituent components, the pathway and kinetics of its assembly have been more difficult to study^24^. Here we have presented rapid AFM imaging data that track MAC assembly at the single-pore level in real-time. Furthermore, we have derived kinetic models for initiation and oligomerization of C9 that explain how rate-limiting assembly intermediates can be captured by our body’s self-defence mechanism to prevent disease.

Based on the results presented here, we define a kinetic pathway of MAC assembly (***Fig. 5***). C5b6 binds to bacterial lipids and serves as a platform for coordinating the sequential assembly of C7 and C8 at the target membrane. This newly formed C5b-8 initiator complex is explicitly required for membrane insertion of the initial C9 molecules. Our data reveal that this initiation phase, which relates to the insertion of the first C9, is the rate-limiting step. A rapid oligomerization phase completes the transmembrane MAC pore, as further copies of C9 bind and insert into the membrane directly from solution. Specifically, the binding of the initial C9 to C5b-8 is characterized by a rate constant that is more than two orders of magnitude smaller than that for the subsequent binding of C9 to C5b-8C9_n_ (1 ≤ *n* < 18), and occurs more slowly than C5b-8 formation. In summary, we show that once activated, the MAC pore rapidly assembles in the target membrane.

**Figure 5:**
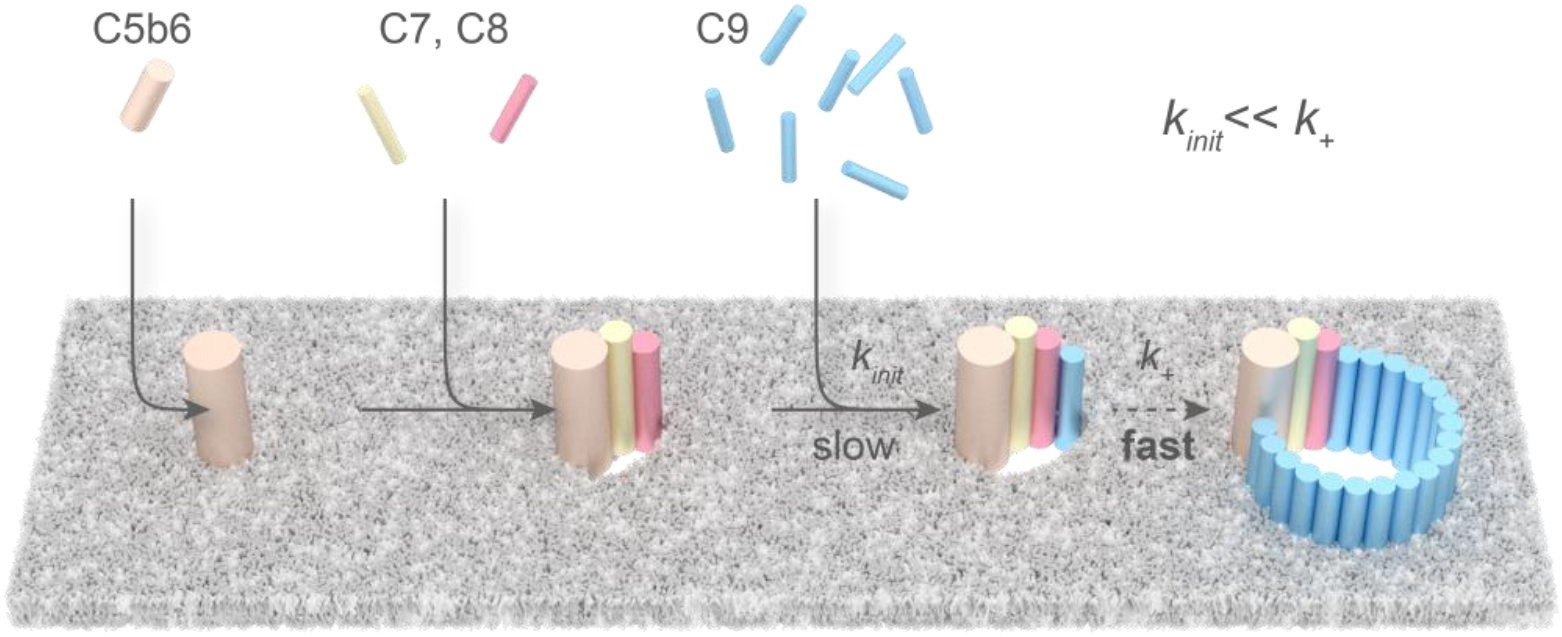
Schematic of MAC assembly. Upon formation, C5b6 binds to the bacterial membrane and recruits C7 and C8 to the nascent MAC. Insertion of the first C9 is the rate limiting step; once this barrier has been overcome, subsequent C9 binding/insertion occurs more than two orders of magnitude faster than the initial C9 insertion.

The transition from soluble monomers to a transmembrane β-barrel pore is accompanied by dramatic conformational changes in the pore-forming membrane attack complex/ perforin/cholesterol dependent cytolysin (MACPF/CDC) domain of complement proteins. The MACPF/CDC domain is comprised of a central kinked β-sheet with two helical sub-regions (TMH; transmembrane hairpins) that unfurl to form membrane-inserted β-hairpins. Interestingly, the rate-limiting step of MAC assembly, as identified here, coincides with unfurling of the first C9’s hairpins into the membrane. This can be related to the recent structural insight that C9 can bind to the C5b-8 initiator complex before inserting into the membrane; however, undergoing the helix-to-hairpin transition is required to propagate oligomerization^15^.

By having distinct initiation and propagation stages, C9 assembly in the MAC resembles pore formation of the related immune protein perforin, where the rate limiting step is the insertion of a small (membrane-bound but not yet membrane-inserted) pre-pore assembly, acting as a nucleation site from which to grow a transmembrane pore ^33^. Although we do not observe any membrane-bound C9 pre-pore assemblies, the C5b-8 initiator complex, together with a C9 molecule yet to undergo its transmembrane transition, could serve a similar function. Our data provide the first experimental evidence that MAC is a growing pore, similar to perforin. These mammalian immune pores differ from bacterial pore-forming proteins from the same superfamily that undergo a concerted oligomeric pre-pore-to-pore transition, after which no further assembly events have been observed^3,28,34^. In a bacterial membrane that harbours a dense proteinaceous network of porins^35^, such a collective pre-pore-to-pore transition is likely to face a large free energy barrier; instead, a growing pore that directly recruits individual monomers from solution can act as a jack that prises open a hole within the porin lattice.

Our results show that C5b6 initiates MAC assembly at the membrane and all subsequent protein components are integrated directly from solution. The C5b6 crystal structure demonstrated that prior to interacting with lipids, C6 membrane-interacting residues remain in their soluble helical form^17^; however, the complex has been shown to associate with lipid bilayers^16^,^32^. It is likely that C5b6 membrane-binding is dominated by electrostatic interactions of negatively charged lipid headgroups (such as PG lipids commonly found in Gram-negative bacteria^36^) with either the thrombospondin (TS)1 domain of C6^37^ or by its unfurled membrane interacting β-hairpin^16^, both of which expose an interface rich in positive charge. Such lipid headgroup dependence also emerges from studies on MAC binding^29,30^ and complement activation on model membranes^38^. The observed membrane binding of C5b6 is consistent with recent work on bacteria^9^, reporting that the downstream efficiency of MAC formation is greatly enhanced when C5b6 is actively formed by C5 convertases bound to the bacterial surface. Our results here provide the rationale for this observation. We propose that nascent C5b does not leave the bacterial membrane upon its generation by the C5 convertase, but needs to recruit C6, C7 and C8 directly to the target membrane. This facilitates binding and oligomerization of C9, thus generating the functional pores that kill the bacterium.

Although the here discussed lipid dependence suggest some MAC specificity for bacterial targets, C5b8-initiator complexes can deposit and progress to cytolytic pores on host cells if not properly controlled. Therefore, human cells express CD59 on their surface, which disrupt MAC assembly from the point of C5b-8 formation onwards^33,41-43^. Specifically, CD59 inhibits MAC assembly by binding to the TMH β-hairpin on the leading-face of C8α (residues 334-385) ^39^, and to a buried 6 amino acid sequence of C9 (residues 366-371) that is exposed upon binding C5b-8^32^. However, it is not known whether C9 has initiated the insertion of its hairpin into the membrane prior to binding CD59. Our data highlight a rate-limiting step at the association of C9 with the C5b-8 MAC precursor. This allows a maximum temporal window for the mechanism by which human cells are protected from autoimmune attack by the MAC: our model favors a mechanism whereby C9 has not unfurled into the membrane immediately upon binding C5b-8, allowing CD59 to inhibit MAC formation. When CD59-mediated inhibition is overcome by antibody-based drugs such as rituximab that facilitate MAC-induced killing of chronic lymphocytic leukaemia B-cells, cell death follows at the ~100 second time scale^7^. This is consistent with the oligomerization kinetics of a single pore observed here *in vitro*, suggesting that only few pores are sufficient to lyse a B-cell.

In summary, we have determined the pathways and kinetics of assembly for a hetero-oligomeric protein complex by molecular-scale measurements. These assembly kinetics govern how MAC kills bacteria and how our body’s self-defence mechanism prevents membrane damage, which may also be relevant for complement dependent cytotoxicity in cancer immunotherapy^7^. Finally, we anticipate that our findings will guide the interpretation of on-going studies towards an atomistic model of MAC structure.

## Methods

### Materials

*E. coli* lipid extract (total), 1,2-dioleoyl-sn-glycero-3-phosphocholine (DOPC), 1,2-dioleoyl-sn-glycero-3-phosphoethanolamine (DOPE), 1,2-dioleoyl-sn-glycero-3-phospho-(1’-rac-glycerol) (DOPG) and 1-palmitoyl-2-(dipyrrometheneboron difluoride)undecanoyl-sn-glycero-3-phosphocholine (TopFluor^®^ PC) were purchased from Avanti Polar Lipids (Alabama, USA) as a powder and stored at −20 °C prior to use. Complement proteins C5b6, C7, C8 and C9 were purchased as proteins purified from human serum from CompTech (Texas, USA) and stored at −80 °C prior to use. For the rapid AFM imaging experiments, lysis assays, negative-stain EM and FRAP experiments shown here, *E.coli* lipid extract was used. For AFM imaging of intermediates, an equimolar mixture of DOPG:DOPE was used. For QCM-D binding assays, a lipid mixture of DOPC:DOPE:DOPG (47.5:47.5:5 mol %) was used, as formation of continuous bilayers containing high molar ratios of charged lipid (> 30 mol %) were not attainable on the silicon dioxide QCM-D sensors used here^40^.

### Preparation of Lipid Vesicles

Pure lipids were dissolved in chloroform at 10 mg/mL and mixed in solution to give a lipid mixture at a desired molar ratio. The lipid-in-chloroform solution was then dried in a glass vial under a stream of nitrogen gas to give 1 mg of lipid as a thin film. The lipid film was hydrated in buffer (20 mM HEPES, 120 mM NaCl, pH 7.4), vortexed and bath sonicated to give a cloudy lipid suspension. The suspension was then passed through a 50 nm polycarbonate membrane (GE Healthcare Lifesciences) 15 times to yield a clear suspension of small unilamellar vesicles (SUVs). All lipid species used had a gel-to-fluid transition below room temperature, and therefore were assumed to be miscible without heating.

### AFM Sample Preparation

Supported lipid bilayers were formed by injecting 4.5 μL of the SUV suspension to a freshly cleaved mica disk (6 mm diameter) under 18 μL of incubation buffer (20 mM HEPES, 120 mM NaCl, pH 7.4). CaCl_2_ solution (2.5 μL, 100 mM in incubation buffer) was added to give a final calcium concentration of 10 mM; this induces the rupture of the vesicles onto the mica support over an incubation period of approximately 30 minutes. Excess vesicles were then removed from the supernatant by rinsing with 500 μL of incubation buffer, to yield a uniform bilayer free of adsorbed vesicles (as assessed by AFM imaging). All SUVs were incubated at room temperature, above their gel-to-fluid transition temperature.

Endpoint MAC pores were formed by incubating the supported bilayer in a humid chamber at 37 °C and sequentially adding complement proteins C5b6, C7, C8 and C9 at 5 min intervals, with a final 15 min incubation after the addition of C9 prior to initiating the AFM experiment. Final concentrations of complement proteins were approximately 80 nM for C5b6, C7 and C8 and 1.4 μM for C9. Excess soluble protein was removed prior to imaging by washing with 5 x sample volume (25-50 μL) of buffer.

‘Real-time’ MAC samples were formed *in situ* within the AFM liquid chamber. For experiments at 37 °C using a Bruker Dimension FastScan, the complement proteins were injected directly onto the supported lipid bilayer whilst imaging. For rapid imaging experiments performed at 30 °C using home-built AFM instrumentation described previously^41^, the sample chamber was passivated with BSA (0.1 mg/mL in incubation buffer) for 15 minutes and rinsed with 500 μL buffer prior to loading the sample with the supported lipid bilayer. Complement proteins were sequentially injected though channels in the sample chamber to the imaging volume, whilst scanning and in the absence of wash steps, to give a final protein concentration of 80 nM for C5b6, C7 and C8 and 1.4 μM for C9.

### AFM Imaging

AFM imaging was performed in fluid using a Bruker Dimension FastScan (for time-lapse imaging at 37 °C), a Bruker BioScope Resolve (for imaging of MAC intermediates) and a home-built instrument with rapid imaging capabilities^41,42^ (for rapid imaging at 30 °C). Imaging was generally performed in off-resonance tapping / fast force-feedback imaging (Bruker’s PeakForce Tapping) mode where force-distance curves were recorded at either 8 or 32 kHz, with amplitudes of 10-20 nm. With these frequencies, images could be collected at 5-100 seconds/frame. Tapping mode of MAC intermediates was performed whilst scanning bidirectionally (i.e. turn around at the end of each imaging line; no ‘retrace’).

Rapid PeakForce Tapping (with the z scanner driving at 32 kHz) was performed with largely custom built hardware as described in detail elsewhere^41^. Importantly, the AFM head has a sufficiently small laser spot to accommodate miniaturized cantilevers, and the high-speed scanner is flexure-based with a 1.8 μm x 1.8 μm x 2 μm range and ~100 kHz *z* bandwidth. Fast force-distance based imaging modes were implemented by sinusoidally modulating the tip-sample distance at a high rate between 16-32 kHz and recording the resulting deflection signal with a significantly higher sampling rate (512 kHz). The resulting periodic hydrodynamic background was recorded slightly above the surface and subtracted from the deflection in real time. The resulting interaction was a sinusoidal force-distance curve where the maximum force was used for feedback. For practical purposes, highest-quality data were recorded at 30 °C; and next compared with results obtained at the physiological 37 °C (see main text).

Commercial FastScan-D cantilevers (Bruker) were used for all experiments, except for images shown in *Fig. 2*, which used pre-release Fast Tapping probes (Bruker; resonance frequency 140 kHz, spring constant 0.3 N m^-1^). FastScan-D cantilevers have a specified spring constant of 0.25 N m^-1^ with a resonance frequency of 110 kHz in liquid; this exceeds our ramping frequency by at least a factor of 3, sufficient to avoid coupling between the ramping frequency and the cantilever resonance. Cantilevers were rinsed in isopropanol:ethanol (1:1) and plasma cleaned in air prior to use.

### AFM Data Processing

Image analysis was performed using Nanoscope Analysis version 1.80 (Bruker)^27^. Briefly, images were plane levelled and line-by-line flattened with the lipid bilayer as a reference. A Gaussian filter with a full-width half-maximum of 2 pixels (corresponding to 4 nm) was used to smooth out high frequency noise where necessary.

Tracking the evolution of a growing pore, and pore counting, was performed as follows, using MATLAB (MathWorks), and the scripts described were used with Bruker’s MATLAB toolbox: NSMatlabUtilites.

AFM video sequences were loaded into MATLAB, and a 1^st^ order plane background subtraction was applied to each image. A template pore was user-selected from the final image in the sequence. This was used as the template in a 2D cross-correlation analysis, applied to each image in the sequence. If features correlated with the template over a given, normalized threshold value (set as 0.6 here; user-adjusted to optimize recognition), such features were identified as MAC pores. The number of features found in each frame was defined as the pore count. Next, using the coordinates already obtained from the 2D cross-correlation analysis, the coordinates of the appearance and growth of unique pores were tracked, and their coordinates stored into a new array (track). This used two further parameters: (i) A maximum linking distance (typically set at ~ 30 pixels), which defines a pore as being the same unique pore as that detected in the previous frame, only if the coordinates of the feature were within the maximum linking distance. (ii) A maximum gap closing (in frames), which defines the number of frames in which a feature (that is within the maximum linking distance) cannot be found and yet is still defined as belonging to the same track (this reduces the risk of artefacts due to image noise). The pore count and coordinates for individual tracks were then saved into a data structure.

Next, using the track coordinates, new image sequences for each pore were cropped to within a radial distance of 25 nm from the centre of the feature, and including some extra frames recorded before the first appearance of a given pore. For each cropped image sequence, the average height of each frame in the sequence was calculated and saved into a new array. This analysis was repeated for several data sets, and the data saved into new data structures. Finally, the cropped image sequences and average height arrays from several experiments were concatenated. Each track was inspected by the user and any false positives (caused by having too small a threshold value for the 2D cross-correlation analysis) were removed. A Savitzky-Golay filter was applied to each remaining average height array to reduce the effect of image noise. A sigmoid function of the form 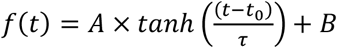 was fitted to each filtered, average height array. A, B, t_0_, and τ are fitting parameters, and 3*τ (which corresponds to ~90% of the transition; see SI Fig. 9) was defined as the width of the transition (and hence reaction time). If, from the fitting, t0 was negative or 3*τ was longer than the video sequence, this was considered a poor fit and the data was removed. For the image sequence tracking a single MAC pore in **fig. 4**, a 1.5 nm Gaussian filter was applied.

### Pore cross-sectional measurements

AFM: Image processing was carried out in Gwyddion^43^. Images were plane levelled and line-by-line flattened with the lipid bilayer as a reference. A Gaussian filter with a full-width half-maximum of 2 pixels was used to smooth out high frequency noise. Following this, a cross-section was then taken diagonally across a single MAC pore (shown in *Fig 1*.) and exported. Data was plotted in Origin.

EM: A MAC pore of structure EM-3134^12^ was imported into Chimera^44^. Volume filtering was performed with *σ* = 14 pixels to obtain a structure with resolution comparable to that of the AFM data. Given a resolution of 8.5 Å for EM-3134, this corresponds to a physical width (of the filter) of 11.9 nm. The volume viewer tool was optimised to ensure the full surface as shown (level 0.002). The model was then coloured by height with a black to white gradient (low to high) of −100 to +200 Å in steps of 75 Å. A top view image was then exported into Gwyddion^43^ for cross-sectional analysis. A cross-sectional measurement was taken horizontally across the pore diameter against the approximate height of a surrounding membrane (50 Å), including the stalk, and exported. Data was plotted in Origin.

### QCM-D Measurements

Quartz crystal microbalance with dissipation monitoring (QCM-D) allows semi-quantification of mass deposition to a quartz crystal sensor. Binding assays were performed by flowing complement proteins over a lipid bilayer supported by the silica-coated QCM-D sensor. Biomolecules interacting with the sensor interface give rise to a change in resonance frequency (Δ*f*) and energy dissipation (Δ*D*) of the quartz sensor. Briefly, a decrease in resonance frequency is proportional to an increase in mass, whilst an increase in dissipation qualitatively correlates with an increase in the ‘softness’ of the film. QCM-D measurements were performed in flow mode at a flow rate of 10 μL/min using a Q-Sense E4 system equipped with four Q-Sense Flow Modules (Biolin Scientific, Vastra Frolunda, Sweden) with a working temperature of 20 °C. Silica coated QCM-D sensors (QSX 303, Biolin Scientific) were used as substrates for supported lipid bilayers. Before injection, C5b6, C7, C8 and C9 were diluted to concentrations 10, 5, 5 and 50 μg/mL (35, 54, 33, 704 nM) respectively in incubation buffer (roughly equivalent to half those use in AFM experiments). Overtones *j* = 3, 5, 7, 9, 11, and 13 were recorded in addition to the fundamental resonance frequency (4.95 MHz). Changes in dissipation (ΔD) and normalized frequency, Δf = Δ*f_j_*/*j*, for *j* = 5 are presented here; all other overtones provided equivalent information.

### Vesicle Lysis Assays

*E. coli* lipid extract was suspended at 7.5 mg/mL in calcein solution (50 mM calcein, 150 mM NaCl, 20 mM Hepes pH 7.4), freeze-thawed 6 times (liquid nitrogen – 65 °C) and extruded through a 100 nm polycarbonate membrane (Whatman) to form unilamellar calcein-encapsulated liposomes. Non-encapsulated calcein was removed through liposomes purification on a gravity-flow Sephadex-G50 (GE Healthcare) column (500 mM Sucrose, 150 mM NaCl, 20 mM Hepes pH 7.4) and liposomes were used immediately. MAC lysis assays of liposomes were performed by sequential addition of C5b6 (5 min, 37 °C), C7 (5 min, 37 °C), C8 and C9 at a mass ratio of 1:1:1:1. In control conditions, identical volumes of protein buffer (120 mM NaCl, 10 mM Hepes pH 7.4) were added instead of protein. Self-quenched encapsulated calcein was un-quenched through its release in the extra-liposomal solution following MAC lesions. Fluorescence was recorded immediately following C9 addition and every minute for 60 minutes on a SpectraMax M2 fluorometer (dual monochromator, ex: 490 nm, em: 520 nm) (Molecular Devices). Background fluorescence was measured from calcein-encapsulated liposomes in the absence of protein and subtracted from the data. To determine the percentage of lysis, the fluorescence was then normalized to the maximal lysis fluorescence estimated after a freeze-thaw cycle of liposomes incubated in 0.25 % sodium dodecyl sulfate (SDS). Fluorescence measures of lysis and controls were always performed on the same batch of liposomes and in three independent replicates.

### Negative-stain EM

Supported lipid bilayers were formed as described for equivalent supported lipid bilayers as used in AFM experiments, using 8 nm thick PELCO^®^ silicon dioxide support films for transmission electron microscopy grids (Agar) as the support instead of mica. Briefly, the glow-discharged grids were incubated with an SUV suspension in calcium containing incubation buffer (20 mM HEPES, 120 mM NaCl, 10 mM CaCl2), rinsed and incubated with complement proteins as described above. Importantly, this allowed us to remove all soluble protein and excess lipid from incubation buffer prior to staining. Samples grids were rinsed with 500 μl incubation buffer, taking care that they remained hydrated throughout, and subsequently stained with 2 % wt/wt uranyl acetate. The sample was incubated with uranyl acetate for 60 seconds and carefully blotted dry, ensuring that the strain was quickly removed to avoid crystallisation of excess uranyl acetate at the surface. Samples were imaged on a Tecnai T12 thermionic filament microscope (Thermo Fisher Scientific) at 120 kV. Images were taken with a defocus of 0.5-1 μm on a Gatan 4k x 4k CCD camera, giving a final pixel size of 1.64 Å.

### Fluorescence Recovery After Photobleaching

*E. coli* lipid extract was doped with 0.5 mol % Topfluor PC by mixing the respective lipids in chloroform and preparing SUVs^27^. A thin mica slide was suspended over a 10 mm glass window in a glass bottomed cell culture plate. Deposition of fluorescently doped SUVs was achieved as described above, taking care to ensure that the mica support remained hydrated at all time. Lipid bilayers were transferred to a FV1200 confocal microscope equipped with a 100x/1.40 oil immersion objective (both Olympus) and a TC-324B automatic temperature controller (Warner Instruments) set to 37 °C. The microscope was further set to a FV10-LD473 473 nm and 15 mW laser diode powered by a FV10-MCPSU power supply, and the Alexa 488 excitation filter and BA490-590 emission filter sets (all Olympus). The fluorescent bilayers were imaged at 2.5 % laser output power across 49.8 μm wide areas and an acquisition speed of 1.64 s per image. FRAP was performed on a circular area of 16.4 μm diameter at the centre of an image for 2 s at a laser output of 80 %. Analysis of the FRAP data was performed according to the Soumpasis model for diffusion limited recovery^24,42^.

### Kinetic Analysis

Given that the initiation and prolongation of C9 binding to C5b-8 occur at such different time scales (*τ*_init_ and *τ*_olig_ or *τ*_+_, respectively), we approximate the reaction kinetics by considering the separate reactions 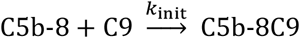 and 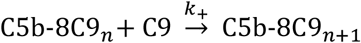 (1 ≤ *n* < 18), while assuming an excess (and therefore approximately constant) amount of C9 in solution (see main text).

The kinetics of the initiation reaction then follows from the differential equation d[C5b-8C9]/d*t* = *k*_init_[C5b-8][C9], where square brackets denote concentrations. For a single C5b-8 complex, we define the initiation probability *p*_init_ ≡ [C5b-8C9]/([C5b-8] + [C5b-8C9]). This leads to a solution of the form *p*_init_ = 1 – exp(−*k*_init_[C9]*t*), such that *k*_init_ ≈ *τ*_init_^-1^[C9]^-1^, with *τ*_init_ determined as illustrated in ***Fig. 3b.***

Defining 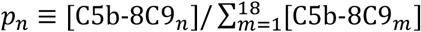, we can describe the subsequent C9 oligomerization via the coupled differential equations d*p*_1_/d*t* = −*k*_+_[C9]*p*_1_, d*p_n_*/d*t* = *k*_+_[C9]*p*_*n*−1_, – *k*_+_[C9]*p_n_* for 2 ≤ *n* < 18, and d*p*_18_/d*t* = *k*_+_[C9]*p*_17_. The average oligomerization rate is given in number of added C9 molecules per unit of time, as 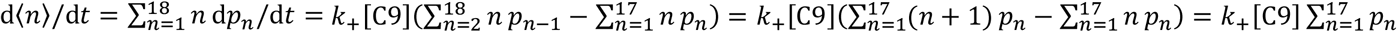. Taking into account that 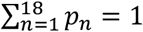 and assuming that *p*_18_ is significantly smaller than 1 (i.e., ignoring the oligomerization of the last few C9s), we find a constant oligomerization rate d〈*n*〉/d*t* ≈ *k*_+_[C9], such that *k*_+_ ≈ *τ*_+_^-1^[C9]^-1^, with *τ*_+_ determined as in ***Fig. 4.***

## Acknowledgements

We thank Richard Thorogate for technical support, Natalya Lukoyanova and Shu Chen for training and advice on EM, and Andrea Slade and James Shaw (Bruker) for advice and assistance with and access to their AFM equipment. This works has been funded by the UK BBSRC and MRC project grants (BB/N015487/1 and MR/R000328/1, to B.W.H.); UK EPSRC and MRC fellowships (EP/M507970/1 to E.S.P.; EP/M506448/1 and MR/R024871/1 to A.L.B.P.); and UK EPSRC investment in AFM equipment (EP/M028100/1). AM and DB are supported by a CRUK Career Establishment Award (C26409/A16099) to D.B. G.E.F and A.P.N acknowledge funding from the European Union FP7/2007-2013/ERC under Grant Agreement No. 307338-NaMic and the European Union H2020 Framework Programme for Research & Innovation (2014-2020); ERC-2017-CoG; InCell; Project number 773091.

## Author contributions

E.S.P. conceived the study, carried out AFM, QCM-D, EM and FRAP experiments, analysed data, led the research and wrote the manuscript. G.J.S. developed and performed tracking analysis of AFM data. A.L.B.P. contributed protocols for, advised on and assisted with AFM experiments, analysed data and wrote the manuscript. A.W.H. performed EM and FRAP experiments. A.P.N. provided home-built instrumentation and assisted with AFM experiments. A.M. performed EM and lysis experiments. A.R.Y. assisted with AFM experiments. A.R. assisted with QCM-D experiments. R.P.R. conceived and advised on QCM-D experiments, and contributed to data analysis and interpretation. G.E.F. provided home-built instrumentation and advised on AFM experiments. D.B. advised on EM and lysis experiments and on structural aspects of the MAC, and wrote the manuscript. B.W.H. conceived the study, developed the tracking analysis of the AFM data, analysed data, led the research and wrote the manuscript. All authors reviewed and commented on the manuscript and its intellectual content.

## Additional information

Supplementary information is available in the online version of the paper. Reprints and permission information is available online. Correspondence and requests for materials should be addressed to E.S.P. and B.W.H.

